# Fungal abundance and community composition associations with environmental and operational factors within and between manufacturing facilities

**DOI:** 10.64898/2026.11.20.768029

**Authors:** Nixon Anthony, Kuan Chu Chen, Padmalya Nisala Supangkat, Wen Chun Chan, Jie Han Li, Li Ya Hung, Kuan Chieh Chen

## Abstract

Fungal communities in manufacturing environments remain understudied despite their potential implications for product quality and occupational health. Surface swabs collected from 19 footwear factories in Vietnam between 2023 and 2025 were used to assess culturable fungal counts, genus-level diversity and community composition. Associations with operational factors and environmental conditions, including temperature, relative humidity (RH), CO₂ and PM₁₀, were examined by separating environmental variation into within-factory and between-factory components. Fungal counts were associated with both operational and environmental factors, with object type showing the strongest operational association. Higher PM₁₀ tended to be associated with lower surface fungal counts within factories, while factories with higher average CO₂ also showed lower counts. *Alternaria* and *Neurospora* were positively associated with within-factory RH, and *Aspergillus* showed a negative between-factory association with CO₂. Community composition varied most strongly among factories, while environmental associations were limited to the combined effects of between-factory conditions at the community level. Overall, environmental indicators showed clearer associations with culturable fungal burden and individual genera than with overall community composition, indicating that fungal counts and community structure capture distinct aspects of contamination in manufacturing environments.

**Importance:** Fungal spores can remain dormant and undetected on factory surfaces until favorable conditions allow germination and growth, by which time products may already be contaminated. Monitoring in manufacturing often relies on colony counts alone. By mapping the count and composition of culturable fungi across footwear factories, this study shows which operational factors and environmental conditions relate to how much fungus is recovered and which genera are present. Several recovered genera include taxa reported in relation to occupant health and material deterioration, although these traits were not assessed here. Combining colony counts with genus identification and factory-level environmental records may help factories prioritize prevention on the surfaces and conditions most associated with fungal presence and abundance.

## Introduction

Fungal communities are an important component of indoor microbiomes, with implications for occupant health (1), material deterioration and product contamination (2, 3). Unlike outdoor environments, indoor spaces are shaped by human activities, building characteristics and environmental management (4). Fungi can enter buildings through outdoor air (5), mechanical ventilation (6), incoming materials and human movement (7, 8), while their persistence depends on their ability to withstand disturbances such as cleaning (9) and fluctuations in environmental conditions (10, 11). Dormant fungal propagules could remain on surfaces until favorable conditions allow germination and proliferate. In manufacturing environments, such processes can contribute to contamination that remains undetected until products exhibit visible fungal growth or deterioration, potentially resulting in substantial product losses (12).

Manufacturing facilities provide a complex setting for examining these interactions. Footwear factories operate as relatively open indoor systems, where fungi may originate from raw materials, personnel, equipment, building surfaces and outdoor air (12). Facilities in tropical regions such as Vietnam also experience warm and humid conditions that may favor fungal persistence (13). Previous studies have examined the distribution of microbial taxa across production flow stages, products and environmental surfaces, which may indicate microbial redistribution within facilities (3). However, it remains unclear whether fungal distributions are more closely associated with production stages or with the functional characteristics of objects (12, 14).

Environmental monitoring is widely used in manufacturing facilities to assess conditions associated with microbial contamination (15). Temperature and relative humidity may influence fungal growth and persistence, and CO₂ and particulate matter (PM₁₀) serve mainly as indirect proxies for occupancy, ventilation and airborne particle dynamics (6). Moreover, variations of local conditions among areas can be described as within factories, while differences between factories may reflect broader building characteristics and operational practices (4, 15). Although indoor microbial communities have been investigated in residential (4), healthcare (16) and food-processing environments (15), comparatively less is known about environmental gradients in open manufacturing facilities. Separating within-factory and between-factory environmental variation may therefore help clarify how local conditions and broader factory differences relate to fungal contamination.

This study characterized culturable fungal count, diversity and community composition on surfaces in 19 footwear manufacturing factories in Vietnam sampled between 2023 and 2025. We addressed two questions: (i) are fungal count, diversity and community composition associated with the same operational factors, including object type, flow stage and factory identity? and (ii) are temperature, relative humidity, CO₂ and PM₁₀ associated with fungal community within and between factories?

## Results

### Operational and environmental associations with fungal count

Sampling was broadly distributed across object types, flow stages and sampling years (Tables S2 and S3), but the share of plates with colony growth varied more among object types (Figure S1). Shoes represented 11.2% of sampled plates but only 6.5% of plates with observed colony growth, while storage and ventilation/dehumidifier surfaces contributed relatively higher proportions of plates with observed colony growth than their sampling proportions. In comparison, distributions across flow stages and years remained broadly similar, although S1 Incoming and 2024 increased in contributions to plates with observed colonies. Mean fungal counts were highest on storage surfaces, followed by building envelopes, ventilation/dehumidifier surfaces and production equipment (Table S2), all of which were classified as environmental surfaces (Table S1). Among product-associated surfaces, raw materials had the highest mean fungal count, whereas finished shoes had the lowest mean count across all object types.

Fungal count was significantly lower on packaging, tables, raw materials, semi-finished products and shoes than on building envelopes (Table 1). Storage, ventilation/dehumidifier and production equipment surfaces did not differ significantly from building envelopes. The fungal count was generally lower from environment surfaces to product surfaces. There were no significant differences among the flow stages. Sampling year effects may reflect unmeasured temporal differences or changes in the factories sampled and were not interpreted further.

**Table 1.** NBMM of fungal count at the plate-grain (n=1,013 plates) for operational predictors. Reference categories were building envelope for object type, S1 Incoming for flow stage, and 2023 for year.

| Predictor | Level | IRR | 95% CI low | 95% CI high | p-value |
|---|---|---|---|---|---|
| Object type | Packaging | 0.42 | 0.28 | 0.63 | **< 0.001** |
| Object type | Production equipment | 0.81 | 0.47 | 1.39 | 0.444 |
| Object type | Raw material | 0.27 | 0.18 | 0.41 | **< 0.001** |
| Object type | Semi-finished | 0.11 | 0.07 | 0.19 | **< 0.001** |
| Object type | Shoe | 0.06 | 0.04 | 0.10 | **< 0.001** |
| Object type | Storage | 1.14 | 0.74 | 1.74 | 0.557 |
| Object type | Table | 0.32 | 0.19 | 0.54 | **< 0.001** |
| Object type | Ventilation/dehumidifier | 1.20 | 0.72 | 1.99 | 0.489 |
| Flow stage | S2 Production | 0.90 | 0.63 | 1.29 | 0.574 |
| Flow stage | S3 Finishing | 0.86 | 0.58 | 1.26 | 0.432 |
| Year | 2024 | 1.59 | 1.05 | 2.41 | **0.028** |
| Year | 2025 | 0.76 | 0.49 | 1.19 | 0.232 |

Indoor CO₂ concentrations varied widely among sampling cells (Table S4), which may reflect differences in occupancy and ventilation (17). Within factories, areas with higher PM₁₀ tended to have lower fungal counts on surfaces, though it did not remain significant after false discovery rate (FDR) correction (Table 2). This pattern may reflect differences between airborne particle concentrations and their deposition onto surfaces, although deposition was not directly measured. CO₂, by contrast, was not related to count within factories, but factories with higher average CO₂ had lower counts. As indoor CO₂ concentrations can vary with occupancy, mechanical ventilation, and operating conditions (15), this association may reflect differences in operating environments across factories. Within- and between-factory slopes differed significantly for CO₂ (Table 2). Temperature and relative humidity showed no significant contrasts. Overall, fungal count varied most with object type. Environmental variables related to count differently within and between factories, separating local conditions from factory-level conditions.

**Table 2.** NBMM of fungal count at EnvCell (n = 245 cells), assessing within- and between-factory components. Estimates are incidence rate ratios (IRR) with 95% CI in parentheses. IRR < 1 indicates a negative association, whereas IRR > 1 indicates a positive association. p-values are followed by FDR-adjusted p-values (q values) in parentheses. Bold indicates p or q < 0.05.

| Variable | within (95% CI) | p within | between (95% CI) | p between | p within vs between |
|---|---|---|---|---|---|
| Temperature | 0.89 (0.73–1.08) | 0.244 (0.488) | 1.39 (0.87–2.24) | 0.171 (0.227) | 0.086 (0.086) |
| RH | 1.00 (0.79–1.27) | 0.983 (0.983) | 0.58 (0.33–1.02) | 0.057 (0.114) | 0.076 (0.086) |
| CO<sub>2</sub> | 1.08 (0.82–1.41) | 0.600 (0.800) | 0.56 (0.36–0.87) | **0.010 (0.041)** | **0.009 (0.035)** |
| PM<sub>10</sub> | 0.72 (0.56–0.94) | **0.013** (0.053) | 1.21 (0.82–1.77) | 0.335 (0.335) | **0.028** (0.057) |

### Operational and environmental associations with diversity

*Aspergillus*, *Penicillium*, and unidentified colonies were the dominant groups across object types (Figure S2), although their relative abundances varied among object types (Figure 1). In contrast, *Cladosporium* showed its highest relative abundances on environmental surfaces, particularly tables and production equipment. *Curvularia* (4.91%) and *Fusarium* (2.31%) were also most abundant on tables (Table S5). *Lichtheimia* reached its highest relative abundance on production equipment (5.69%), whereas *Trichoderma* was most abundant on semi-finished products (4.88%) and raw materials (4.51%). *Neurospora* was distributed across several object types, with relatively higher abundances on semi-finished products (3.13%), raw materials (3.11%), ventilation/dehumidifier surfaces (3.07%), storage surfaces (2.61%), and shoes (2.32%). *Paecilomyces* remained relatively rare and was detected only on raw materials, tables, packaging, production equipment, and building envelopes.

**Figure 1.**
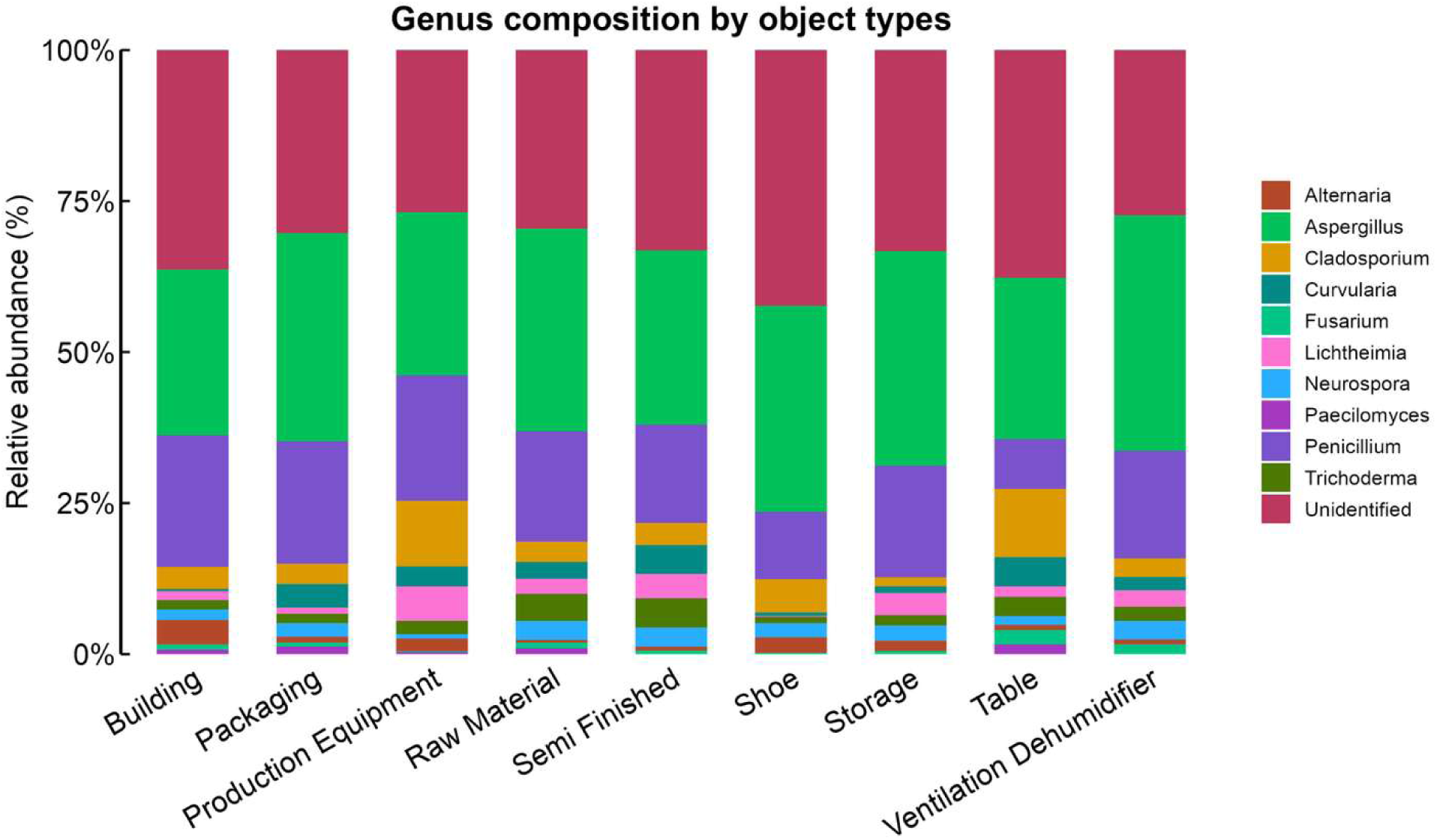
Proportional analysis percentages of genus across object types.

Hill diversity did not vary among flow stages, but all three diversity orders varied among object types (Table 3), so object types contained different numbers of genera and different shares of common and dominant genera. Environmental effects on diversity were generally weak, although factories with higher average PM₁₀ tended to have lower diversity at all three Hill orders (Figure 2; Table S6).

**Figure 2.**
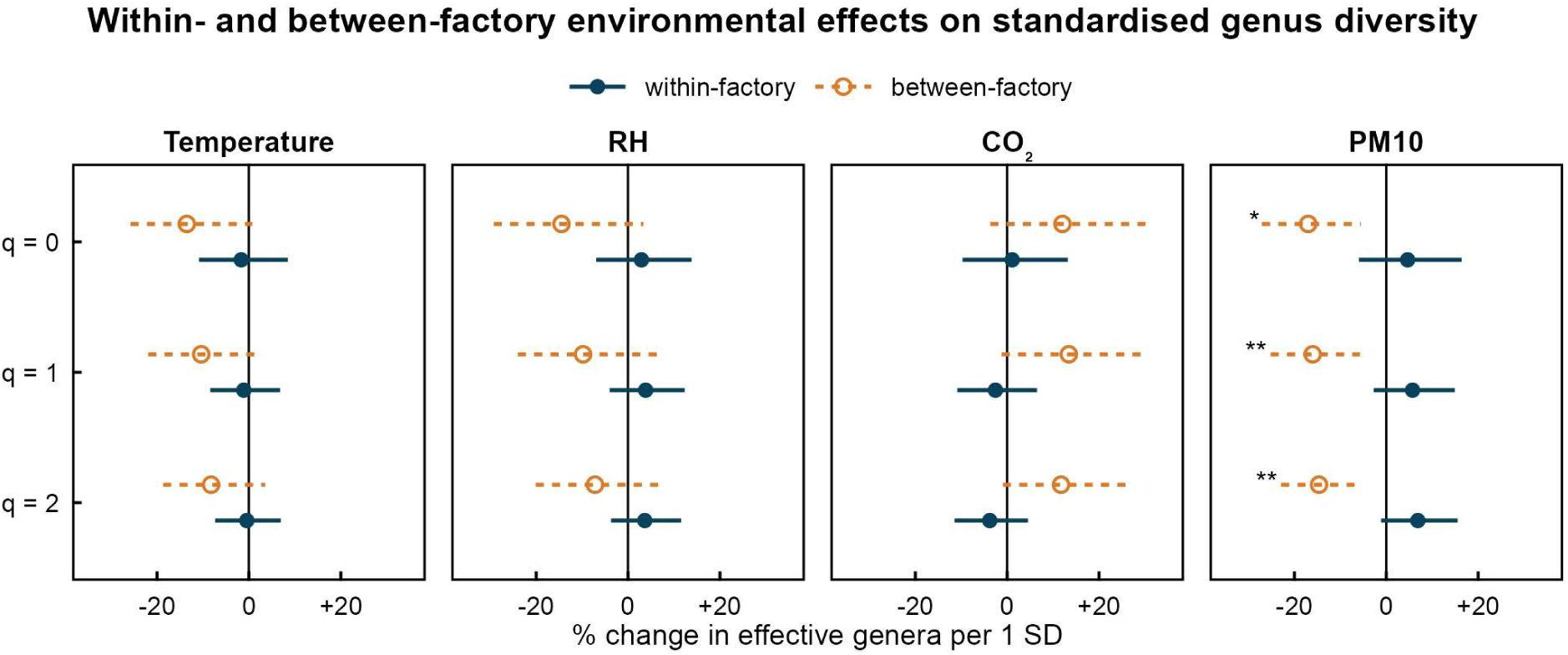
Within- and between-factory environmental effects on coverage-standardized genus-level Hill diversity (n=206 cells). Points represent estimated percentage changes in effective genera associated with a 1-SD increase in parameters. Within-factory effects represent local environmental deviations, whereas between-factory effects represent differences in average environmental conditions among factories. Asterisks indicate significance *p* < 0.05.

**Table 3.** Global associations (*p*-value) of operational factors with coverage-standardized genus-level alpha diversity (n=546 plates).

| Predictor | Richness (q = 0) | Shannon (q = 1) | Simpson (q = 2) |
|---|---|---|---|
| Object type | **0.001** | **0.004** | **0.007** |
| Flow stage | 0.770 | 0.990 | 0.940 |
| Sampling year | **0.002** | **0.015** | 0.120 |

### Multivariate analysis

Operational factors related differently to genus abundance and composition (Table 4), and sampling year affected the most genera. Object type was significantly associated with the abundance of *Aspergillus*, *Lichtheimia*, *Neurospora*, and *Penicillium*, whereas flow stage showed limited associations, with only *Trichoderma* abundance remaining significant after FDR correction. No object-type or flow-stage effect on genus composition remained significant after correction, so these factors changed how much of each genus was recovered more than their relative proportions. Global multivariate tests detected only environmental associations in genus abundance or composition (Table S7). Relative humidity showed pooled associations with both abundance (p = 0.039) and composition (p = 0.028), while between-factory PM₁₀ showed an association with composition (p = 0.047). However, none of these associations remained significant after FDR correction.

**Table 4.** Genus-specific associations of operational factors with fungal abundance and community composition at the plate level (n = 663). Values are p-values from drop-one likelihood-ratio tests, with FDR-adjusted p-values (q values) in parentheses. Bold indicates p or q < 0.05.

| Model | Genus | Year | Object | Flow stage |
|---|---|---|---|---|
| Abundance | *Alternaria* | **0.001 (0.002)** | 0.086 (0.155) | 0.114 (0.257) |
|  | *Aspergillus* | **0.002 (0.004)** | **0.001 (0.003)** | 0.260 (0.326) |
|  | *Cladosporium* | **0.007 (0.011)** | 0.302 (0.422) | **0.044** (0.132) |
|  | *Curvularia* | **0.001 (0.002)** | 0.328 (0.422) | 0.290 (0.326) |
|  | *Fusarium* | 0.573 (0.573) | 0.695 (0.695) | **0.020** (0.090) |
|  | *Lichtheimia* | 0.175 (0.225) | **0.001 (0.003)** | 0.243 (0.326) |
|  | *Neurospora* | **0.001 (0.002)** | **0.017 (0.038)** | 0.870 (0.870) |
|  | *Penicillium* | **0.001 (0.002)** | **0.001 (0.003)** | 0.181 (0.326) |
|  | *Trichoderma* | 0.379 (0.426) | 0.488 (0.549) | **0.003 (0.027)** |
| Composition | *Alternaria* | **0.025 (0.032)** | 0.681 (0.766) | 0.468 (0.999) |
|  | *Aspergillus* | 0.082 (0.092) | 0.409 (0.766) | 0.895 (0.999) |
|  | *Cladosporium* | **0.003 (0.011)** | 0.860 (0.860) | **0.037** (0.111) |
|  | *Curvularia* | **0.004 (0.011)** | 0.618 (0.766) | 0.999 (0.999) |
|  | *Fusarium* | **0.021 (0.032)** | 0.251 (0.766) | **0.030** (0.111) |
|  | *Lichtheimia* | **0.007 (0.013)** | **0.007** (0.063) | 0.720 (0.999) |
|  | *Neurospora* | **0.005 (0.011)** | 0.580 (0.766) | 0.820 (0.999) |
|  | *Penicillium* | **0.001 (0.009)** | 0.532 (0.766) | 0.680 (0.999) |
|  | *Trichoderma* | 0.402 (0.402) | 0.342 (0.766) | **0.013** (0.111) |

At the genus level, responses to environmental variables varied among genera, between abundance and composition, and between scales (Figures 3 and 4). In pooled analyses, *Alternaria* and *Neurospora* increased with RH in both abundance and composition, while only *Neurospora* remained significant after FDR correction. *Aspergillus* decreased with CO₂ in abundance models, and *Penicillium* abundance decreased with temperature, but neither remained significant after correction.

**Figure 3.**
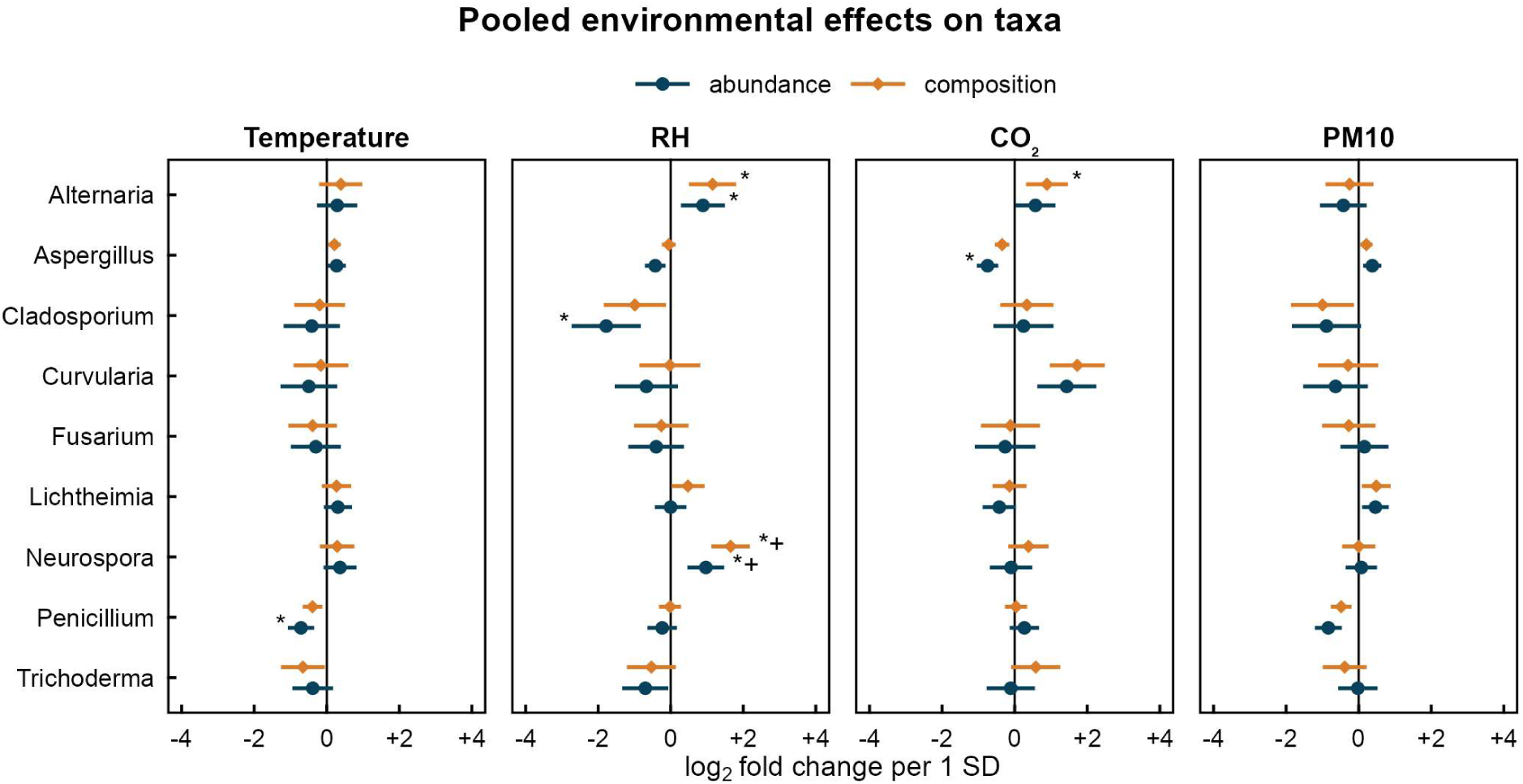
Pooled environmental associations with genus-level fungal abundance and composition at the EnvCell grain (n=221 cells). Points show estimated log₂ fold changes per 1-SD increase in temperature, RH, CO₂, or PM10, with horizontal bars representing 95% confidence intervals. (●) represent abundance models and (◆) represent composition models. Positive values indicate positive associations and negative values indicate negative associations. Asterisks indicate significance *p* < 0.05 and (^+^) indicate associations remaining significant after false-discovery-rate correction (*q* < 0.05).

**Figure 4.**
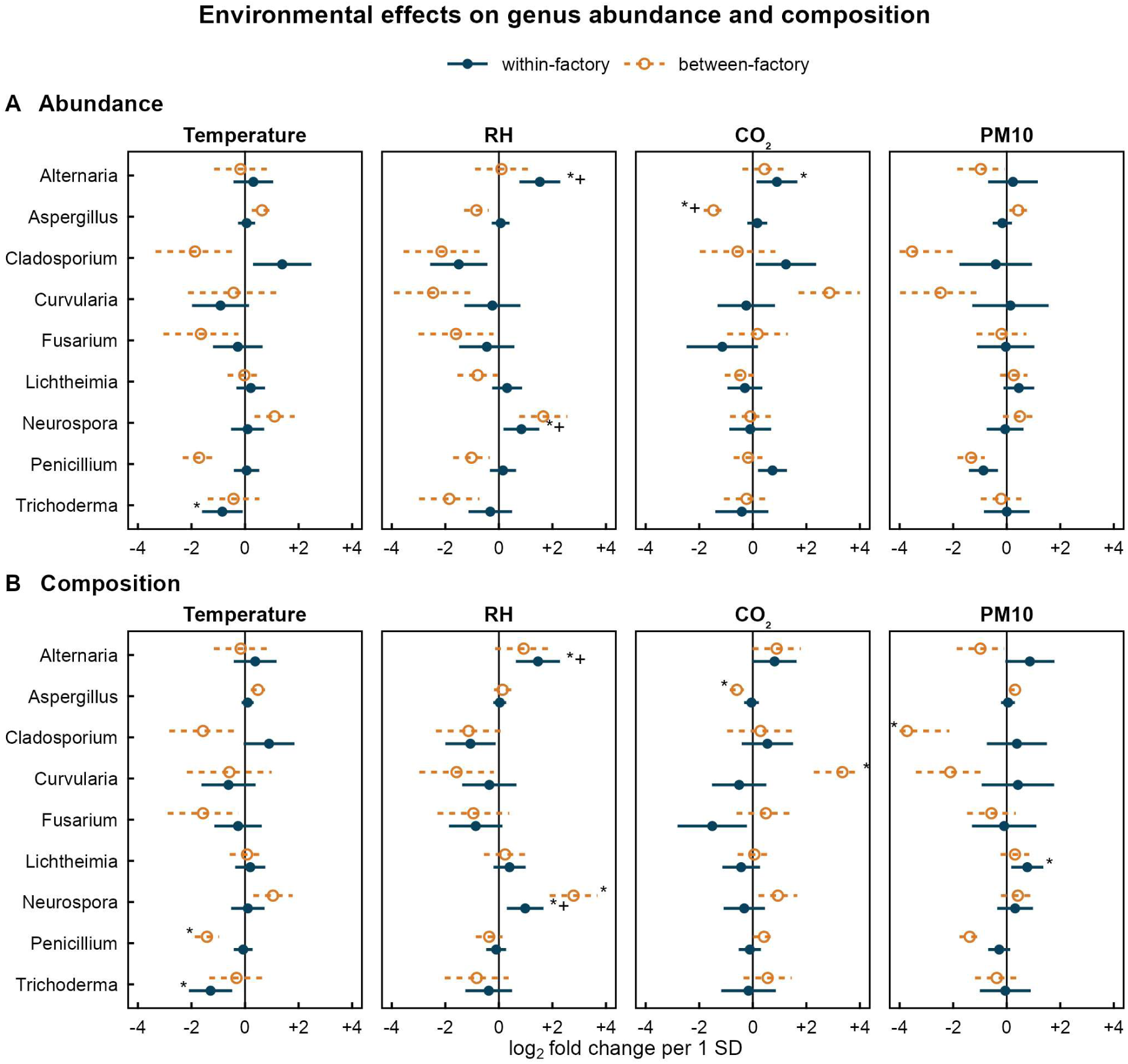
Within-(●) and between-(**○**) factory environmental associations on genus-level fungal abundance and composition at the EnvCell grain (n=221 cells). Points show estimated log₂ fold changes per 1-SD increase in temperature, RH, CO₂, or PM10, with horizontal bars representing 95% confidence intervals. (A) Abundance, (B) Composition. Asterisks indicate significance *p* < 0.05 and (^+^) indicate associations remaining significant after false-discovery-rate correction (*q* < 0.05).

Within factories (Figure 4), *Alternaria* and *Neurospora* increased with RH in both abundance and composition. *Trichoderma* decreased with temperature in both abundance and composition, *Alternaria* abundance increased with CO₂, and the proportion of *Lichtheimia* increased with PM₁₀. Between factories, *Aspergillus* abundance was lower in factories with higher average CO₂. *Penicillium* made up a smaller share of the assemblage in warmer factories, and *Cladosporium* a smaller share in factories with higher PM₁₀, whereas *Curvularia* and *Neurospora* made up larger shares in factories with higher CO₂ and higher RH, respectively.

Overall, object type was linked more to how much of each genus was recovered than to which genera were present. Environmental conditions acted on particular genera depending on their scale.

### Community composition

Variance partitioning showed that factory accounted for the largest share of explained variation in fungal community composition, with environmental predictors contributing additional variation, including fractions shared with factory (Figure S4). Collectively, environmental predictors accounted for 7.61% of Bray–Curtis and 4.74% of Jaccard variation, while flow stage contributed only a small unique fraction to Jaccard variation (0.21%). Shared fractions among operational predictors were limited, and most community variation remained unexplained (77.02% for Bray–Curtis and 80.61% for Jaccard).

In the PCoA, grouping by factory left the least residual variation (Figure S5), and PERMANOVA confirmed that factory accounted for the largest proportion of variation in Bray– Curtis (R² = 17.94%) and Jaccard (R² = 14.57%) distances (Table 5). Sampling year accounted for smaller proportions of variation (2.76% and 2.32%, respectively), followed by object type (1.69% and 1.81%), which remained significant for both distance measures. Flow stage explained little variation and was not significant (0.29% and 0.38%). Dispersion tests detected significant differences among factories, flow stages and sampling years, but not among object types, suggesting that object-type differences in community composition were less likely to be explained by differences in within-group dispersion.

**Table 5.** PERMANOVA and dispersion tests of operational differences (n=666 swabs). The proportion of variation explained by each factor is expressed as R² (%).

| Distance | Variable | R² | p | p (Dispersion) |
|---|---|---|---|---|
| Bray–Curtis | Factory | 17.94 | **0.001** | **0.001** |
| Bray–Curtis | Year | 2.76 | **0.001** | **0.001** |
| Bray–Curtis | Object type | 1.69 | **0.007** | 0.106 |
| Bray–Curtis | Flow stage | 0.29 | 0.253 | **0.036** |
| Jaccard | Factory | 14.57 | **0.001** | **0.001** |
| Jaccard | Year | 2.32 | **0.001** | **0.001** |
| Jaccard | Object type | 1.81 | **0.004** | 0.235 |
| Jaccard | Flow stage | 0.38 | 0.120 | **0.028** |

Environmental variables were more clearly related to composition when assessed jointly than individually. Joint between-factory environmental predictors showed an association with Bray– Curtis composition (R² = 10.91%, p = 0.014; Table S8), whereas individual environmental predictors generally explained smaller proportions of community variation in pooled (Figure 5), within-factory (Figure S6), and between-factory analyses (Figure S7). Environmental conditions therefore related to composition mainly as a combined factory-level signal.

**Figure 5.**
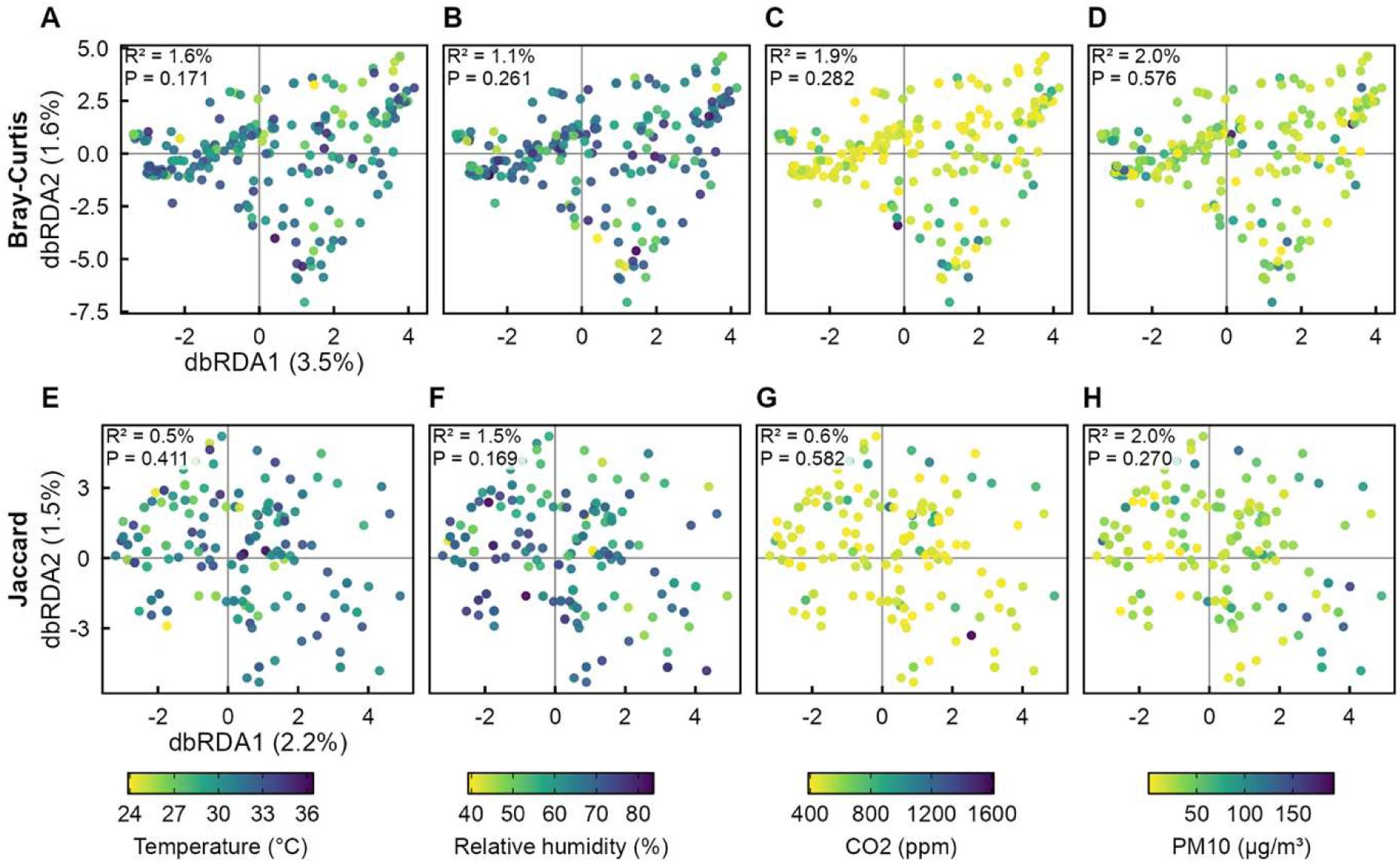
dbRDA of fungal community against environmental variables (n=222 cells). R² values represent the proportion of variation explained by the environmental variables, with residual variation representing the unexplained component.

## Discussion

Object type was the operational factor most strongly associated with fungal count. Object type includes both physicochemical features that influence fungal spore adhesion (18) and operational exposure pathways caused by human handling, cleaning procedures, and dust deposition (12, 19). Fungal counts did not differ among flow stages. Earlier research has shown that microbial transfer occurs easily via surface-touch networks and human interaction (20), resulting in common fungal taxa on environmental surfaces, equipment, and objects (3). These mechanisms are consistent with the broad, shared genus pool observed across object types (12).

Environmental variables were also related to fungal count. Within factories, surfaces in areas with higher PM₁₀ tended to have lower counts, which may reflect particle resuspension from surfaces into air (4) or differences in ventilation operation (5). Between factories, higher counts in factories with lower average CO₂ may reflect greater air exchange and therefore more outdoor spore input (17, 21). Operational factors therefore related to surface fungal count, while environmental variables acted as indirect measures of factory conditions that affect how much culturable fungus is recovered.

Responses to environmental variables also differed among genera and between scales. Previous research has linked *Alternaria* to increased moisture in carpet dust and drywall substrates (22), which fits its positive association with RH within factories. Outdoors, by contrast, dry conditions favor the release of *Alternaria* spores into the air, whereas rainfall washes them out (23). Humidity may therefore act in opposite directions on growth on surfaces and on spore release into air, and surface swabs cannot separate the two. Similarly, *Neurospora* has been found in larger abundances in humid indoor conditions (2). *Penicillium* made up a smaller share of the assemblage in warmer factories, a pattern consistent with its reported higher abundance at higher latitudes (24), which could also support abundance at lower temperature.

Other genus responses may reflect differences in dispersal and indoor air conditions, although the mechanisms remain uncertain (4). Lower *Aspergillus* abundance in factories with higher average CO₂ could reflect differences in ventilation and outdoor air exchange (21). Associations of PM₁₀ in *Cladosporium* and *Lichtheimia* may reflect particle-borne dispersal from outdoors (21) and resuspension of settled particles indoors (8). *Cladosporium* includes important fungal allergens (25), *Curvularia* includes opportunistic pathogens (26), and *Lichtheimia* belongs to the *Mucorales*, a group containing opportunistic human pathogens (27). *Paecilomyces*, although reported in indoor dust and material deterioration studies (28, 29), was rarely recovered in the present study.

Fungal community composition varied more strongly among factories than with measured environmental conditions. Operational practices, mechanical ventilation and occupancy have been associated with indoor community assembly in other facilities (15, 24). Because between-factory environmental terms are factory averages, they represent only part of the factory effect. Environmental conditions related to composition mainly as a combined factory-level signal, whereas individual variables were associated with fungal count and with a few genera. This agrees with studies in which different environmental characteristics determine quantity and composition (24).

Fungal count and community composition responded to different operational factors. Fungal count varied mainly with object type, and composition varied mainly among factories and only slightly among object types. Several recovered genera include taxa relevant to indoor health and material deterioration, although these traits were not assessed here. Combining fungal counts with genus identification may therefore help each factory target operational practices to the surfaces and conditions linked to higher contamination.

Several limitations should be acknowledged. Culture methods capture only fractions of fungal communities and may underestimate overall taxonomic richness (30, 31). Restricting identification to selected target genera further limited the resolution of diversity and community composition analyses. Additionally, contamination sources and potential transfer pathways between environmental surfaces and products were not directly examined. Moreover the absence of detailed operational records, including ventilation systems, cleaning practices, and occupancy, also constrained the interpretation of factory-specific differences. Future studies could combine broader molecular identification with sampling of contaminated or rejected products and detailed operational monitoring to better characterize fungal diversity and trace potential contamination sources.

## Materials and Methods

### Data background

Sampling was conducted by the Mold Research Center, YCM PRODUCTS CO., LTD., at 19 footwear manufacturing factories in Vietnam between 2023 and 2025. Each factory was sampled during daytime working hours in one or more of these years. Surfaces were swabbed following a previously described method (12), yielding 1,013 swab samples, of which 728 showed fungal growth. The sampled surfaces were classified according to the categories listed in Table S1. Swab samples were plated on potato dextrose agar (PDA) supplemented with chloramphenicol and incubated at 30°C for 72 h (12). Environmental parameters, including temperature, relative humidity, CO₂ and PM₁₀, were measured once per report area (12). To avoid pseudoreplication, an environmental sampling unit (EnvCell) was defined by the combination of sampling year, factory, date, warehouse and report area (32, 33). The measurements were aggregated into 245 EnvCells, of which 226 contained samples with fungal growth. Analyses were conducted at the appropriate observational level, with environmental predictors standardized before modeling (34) and decomposed into within- and between-factory components (35).

### DNA and data sequencing

A morphological reference library (Table S9) was developed using fungal isolates recovered from the factory surveys, which are also commonly cultured in indoor environments. Selected isolates were cultured on potato dextrose agar (PDA) for 5 days before submission to an external laboratory for molecular identification (12). Genomic DNA was extracted using the Kaneka Easy DNA Extraction Kit version 2. The internal transcribed spacer (ITS) region was amplified using forward primer ITS5 and reverse primer ITS4 (36) while the translation elongation factor 1-alpha (TEF1-α) gene was amplified using forward primer EF-1 and reverse primer EF-2 for *Fusarium* isolates (37). PCR amplification was performed using PowerAmp 2X PCRmix-Green (Bioman), followed by Sanger sequencing using BigDye Terminator v3.1 chemistry on an Applied Biosystems 3730xl DNA Analyzer. The resulting sequences were compared with reference sequences using BLAST, and the best matches, sequence identity, query coverage, and representative colony photograph were recorded using Nikon SMZ745T. Additionally, *Cladosporium oxysporum* (BCRC 30813), obtained from the Bioresource Collection and Research Center, was included as a comparative morphological reference.

### Regression analyses

All statistical analyses were performed in R version 4.5.0. Fungal count was expressed as total colony-forming units (CFU) and analyzed using negative binomial mixed-effects models (NBMMs) with the *nbinom2* parameterization in *glmmTMB* (38). Operational associations were assessed at the plate level (n= 1,013), with object type, sampling year, and flow stage as fixed effects and factory and environmental exposure cell (EnvCell) as random intercepts.

Alpha diversity was quantified using Hill numbers of orders q = 0, 1, and 2 (39), calculated across ten target genera. Unidentified colonies were excluded because they did not represent a single taxon. To account for unequal colony recovery, assemblages were standardized to 96% sample coverage using iNEXT (40, 41) (Figure S3). Assemblages containing fewer than two target colonies were excluded, retaining 546 plates and 206 EnvCells (Figure S3). Log-transformed Hill numbers were analyzed using linear mixed-effects models (*lmerTest*) (42). EnvCell-level models included sampling year, flow stage, and the within- and between-factory components of temperature, relative humidity, CO₂, and PM₁₀ as fixed effects, with factory as a random intercept. Separate models were fitted for each Hill order.

Genus-level abundance and composition were analyzed using multivariate negative binomial generalized linear models (*mvabund::manyglm*) (43). Unidentified colonies were excluded to focus on taxonomically assigned genera, while *Paecilomyces* was excluded because it occurred in fewer than 5% of sampled plates, leaving nine genera for analysis. The retained dataset comprised 663 plates and 221 EnvCells. Abundance represented genus-specific colony counts, whereas composition represented the relative contribution of each genus within the nine assessed genera (44). Plate-level models included factory, sampling year, flow stage, and object type as fixed effects. EnvCell-level models included the within- and between-factory components of the four environmental predictors. Offset terms were defined differently as the logarithm of total colony counts across the nine assessed genera for composition models and the logarithm of the number of plates per EnvCell for cell-level abundance models (44). Predictors were evaluated using drop-one likelihood-ratio tests with 999 resamples, applying PIT-trap resampling of plates at the plate level (45) and whole-factory resampling at the EnvCell level. Genus-specific p-values were adjusted using the Benjamini–Hochberg procedure within each predictor (46).

### Community analysis

Fungal community composition was assessed using Bray–Curtis and Jaccard dissimilarities (vegan::vegdist) (47). Differences among factories, object types, flow stages, and sampling years were evaluated using PERMANOVA (vegan::adonis2) with 999 permutations (n=666 swabs), while homogeneity of multivariate dispersion was assessed using PERMDISP (vegan::betadisper). Variation partitioning (vegan::varpart) quantified the unique and shared contributions of operational and environmental predictors using adjusted R² (48), and principal coordinate analysis (PCoA; stats::cmdscale) was used for visualization. Environmental associations with community composition were further examined at the EnvCell level (n=222 cells) using distance-based redundancy analysis (vegan::dbrda), controlling for sampling year. Environmental predictors were evaluated collectively and separately across pooled, within-factory, and between-factory conditions. Pooled analyses used the environmental measurements, whereas within- and between-factory analyses accounted for environmental variation at each scale while adjusting for the corresponding components at the other scale. Significance was assessed using 999 permutations, with EnvCells permuted within factories for within-factory tests and whole factories permuted for between-factory tests.

## Acknowledgments

We thank Tri-I Biotech Inc. for sequencing support and the personnel of the YCM Mold Research Center for their assistance during this work. During the preparation of this work, the authors used ChatGPT to improve language and readability, and Claude for code development and data analysis. The authors reviewed and edited all output and take full responsibility for the content of the publication.

## Author contribution

Nixon Anthony (Data curation, Formal analysis, Project administration, Visualization, Writing – original draft), Kuan Chu Chen (Investigation, Validation, Writing – review & editing), Padmalya Nisala Supangkat (Validation, Writing – review & editing), Wen Chun Chan (Conceptualization, Methodology, Supervision), Jie Han Li (Conceptualization, Methodology), Li Ya Hung (Investigation) and Kuan Chieh Chen (Funding acquisition, Resources).

## Data availability

Sequencing and reference data are provided in the supplementary material. The analysis code is available upon reasonable request from the Mold Research Center, YCM PRODUCTS CO., LTD. (contact corresponding author).

---

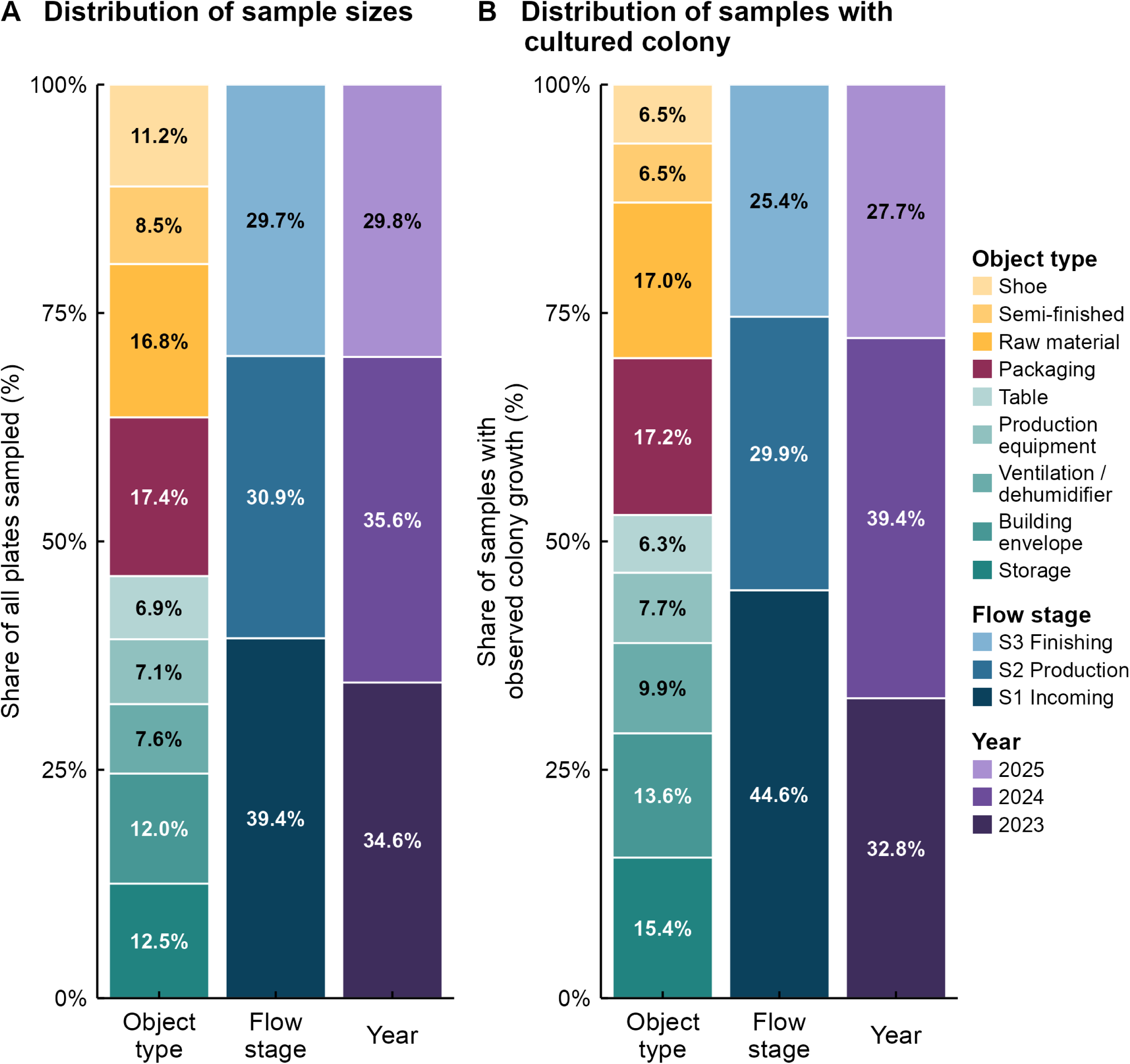

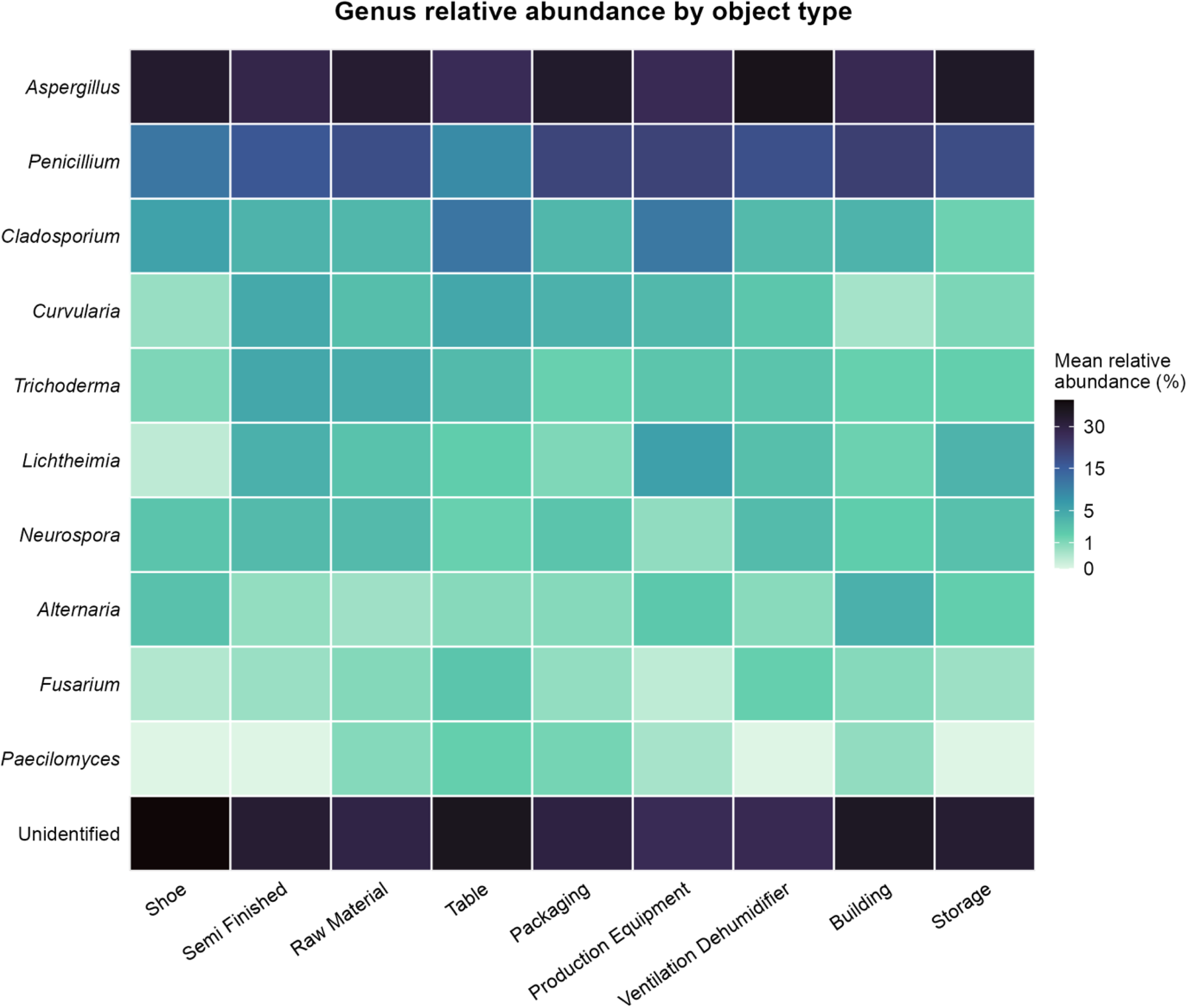

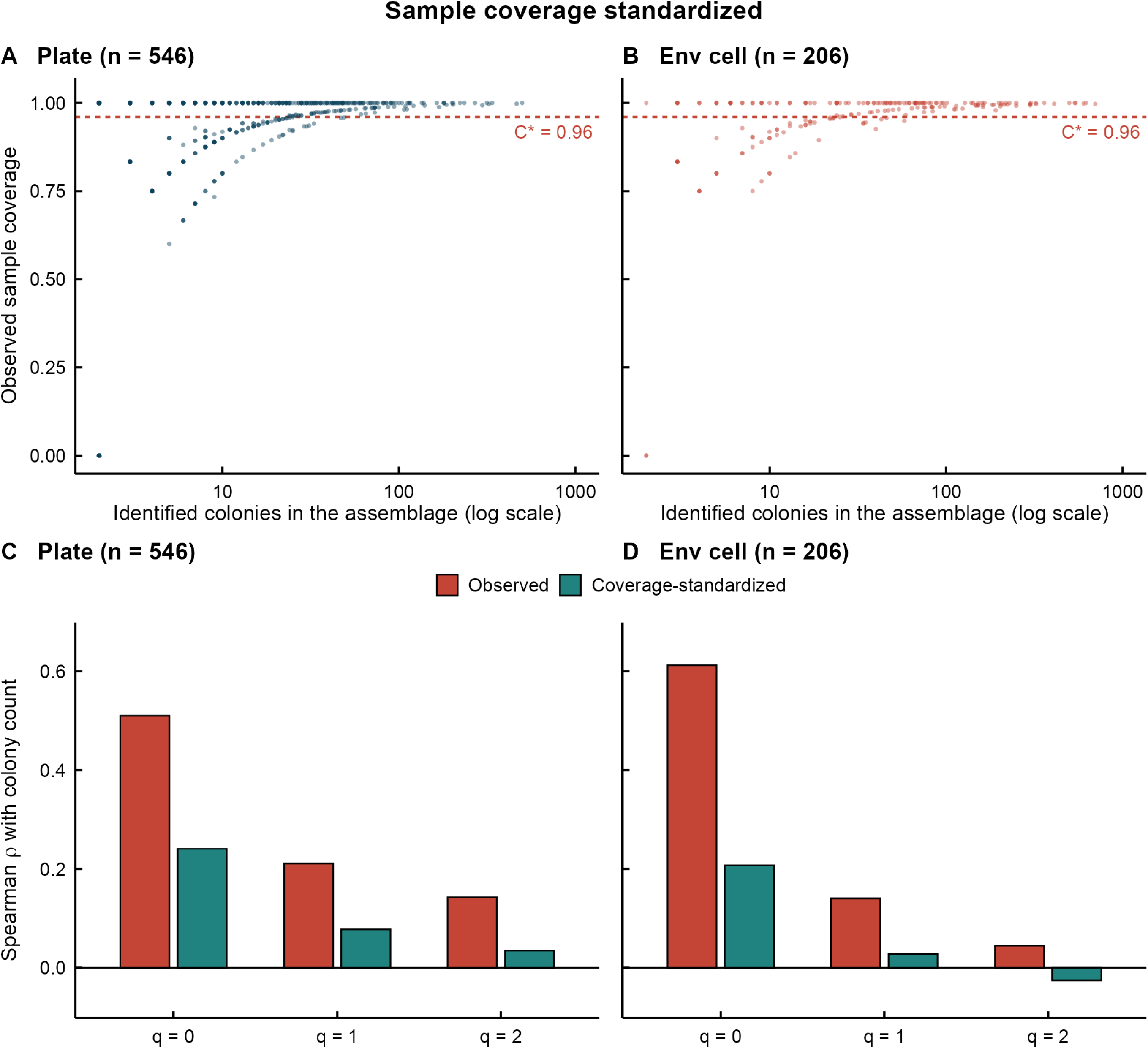

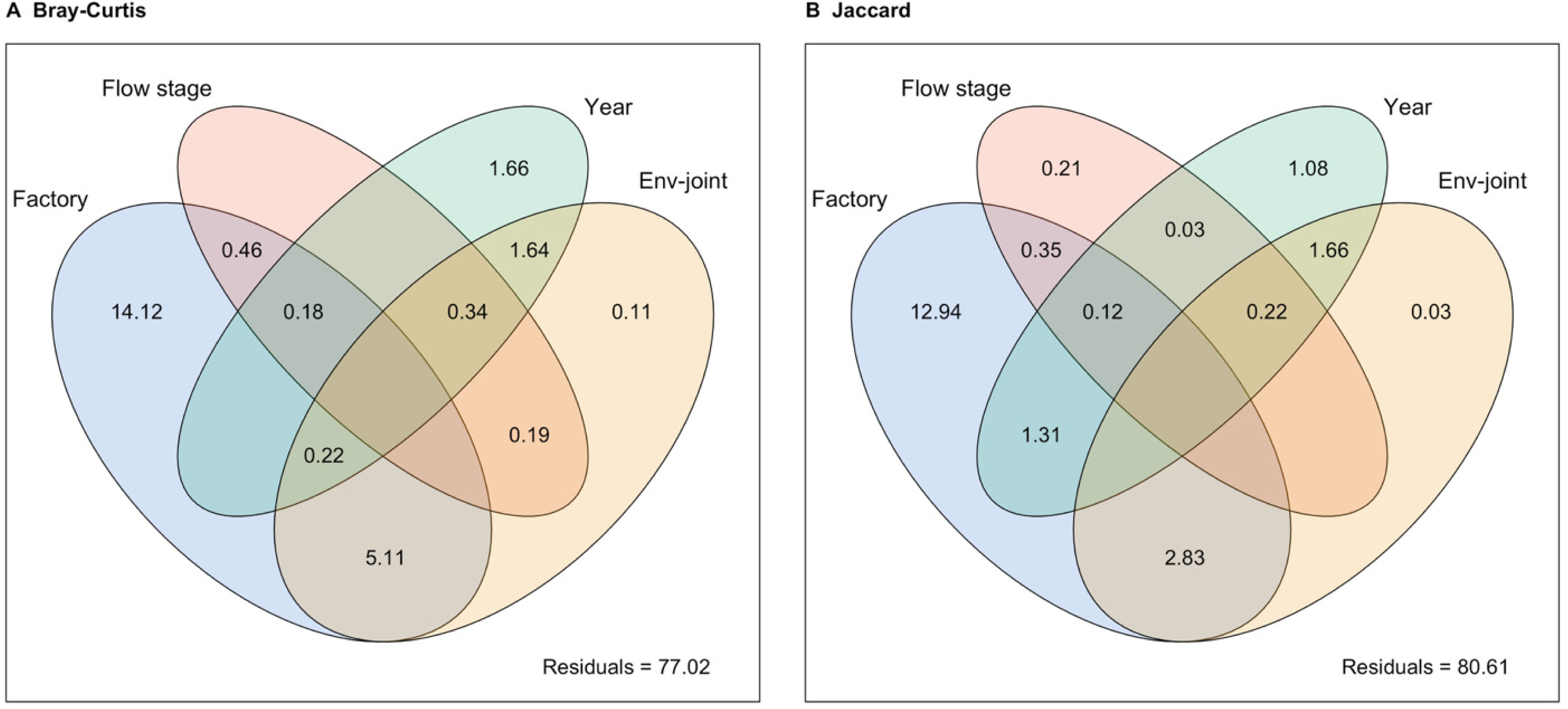

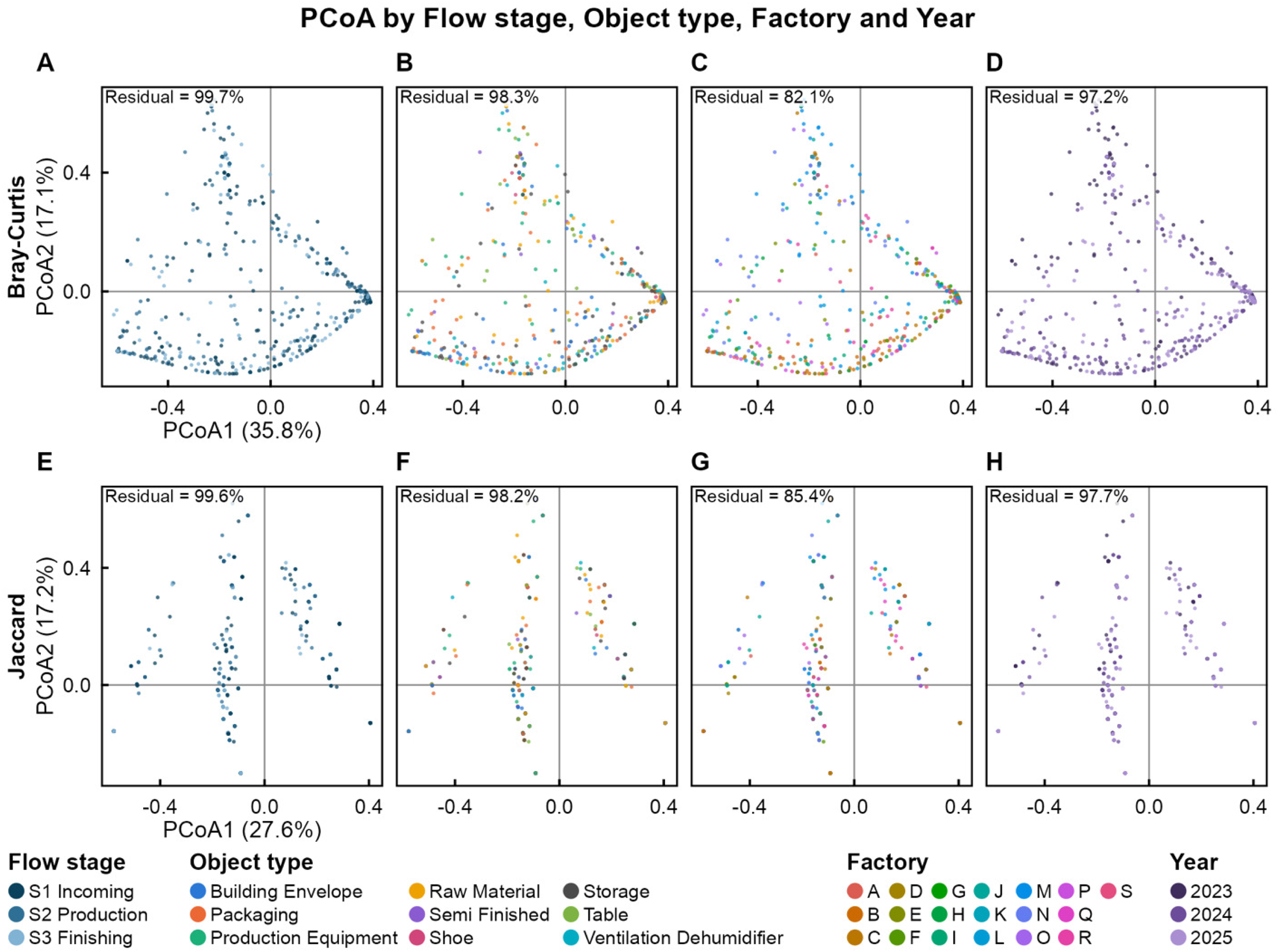

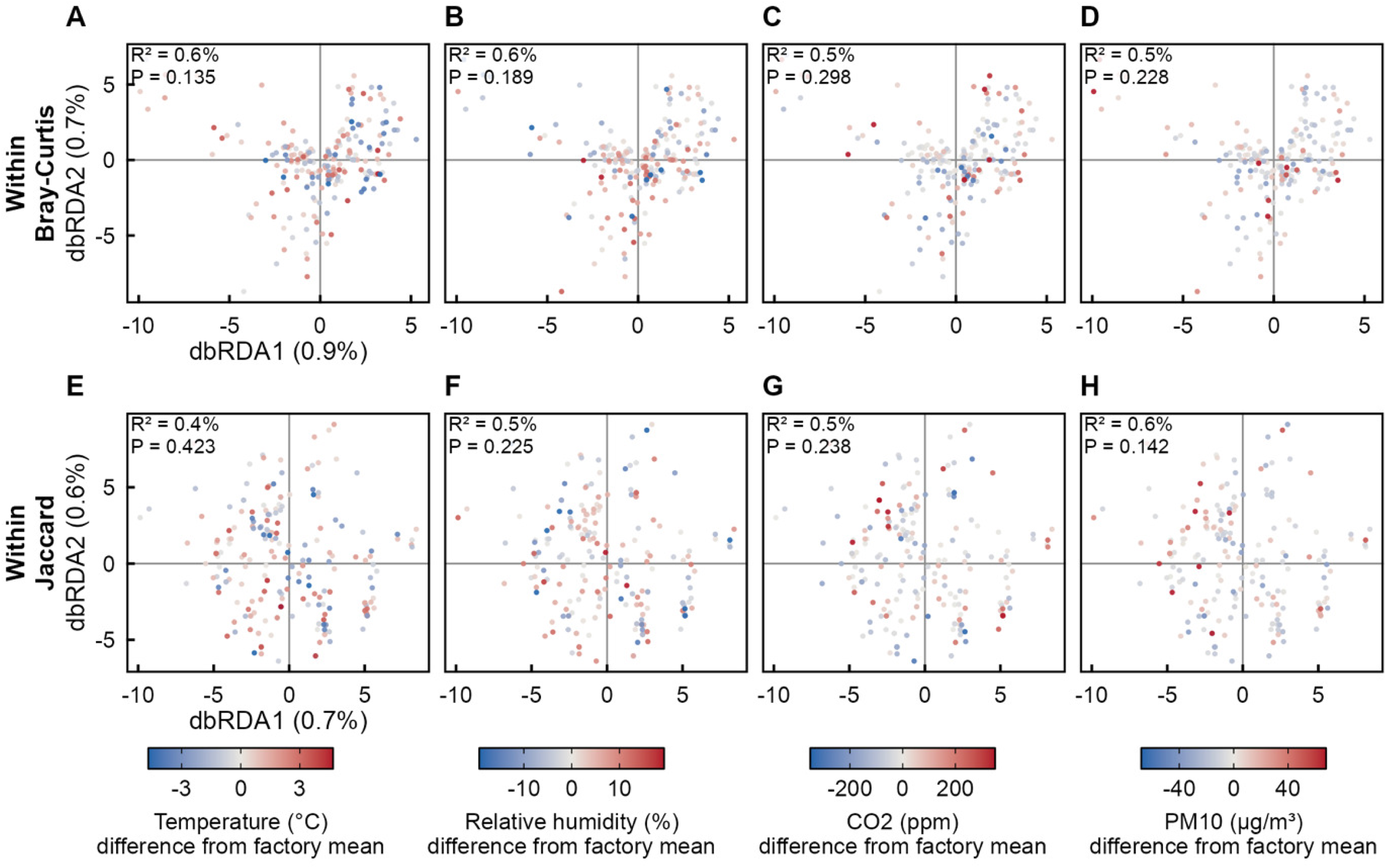

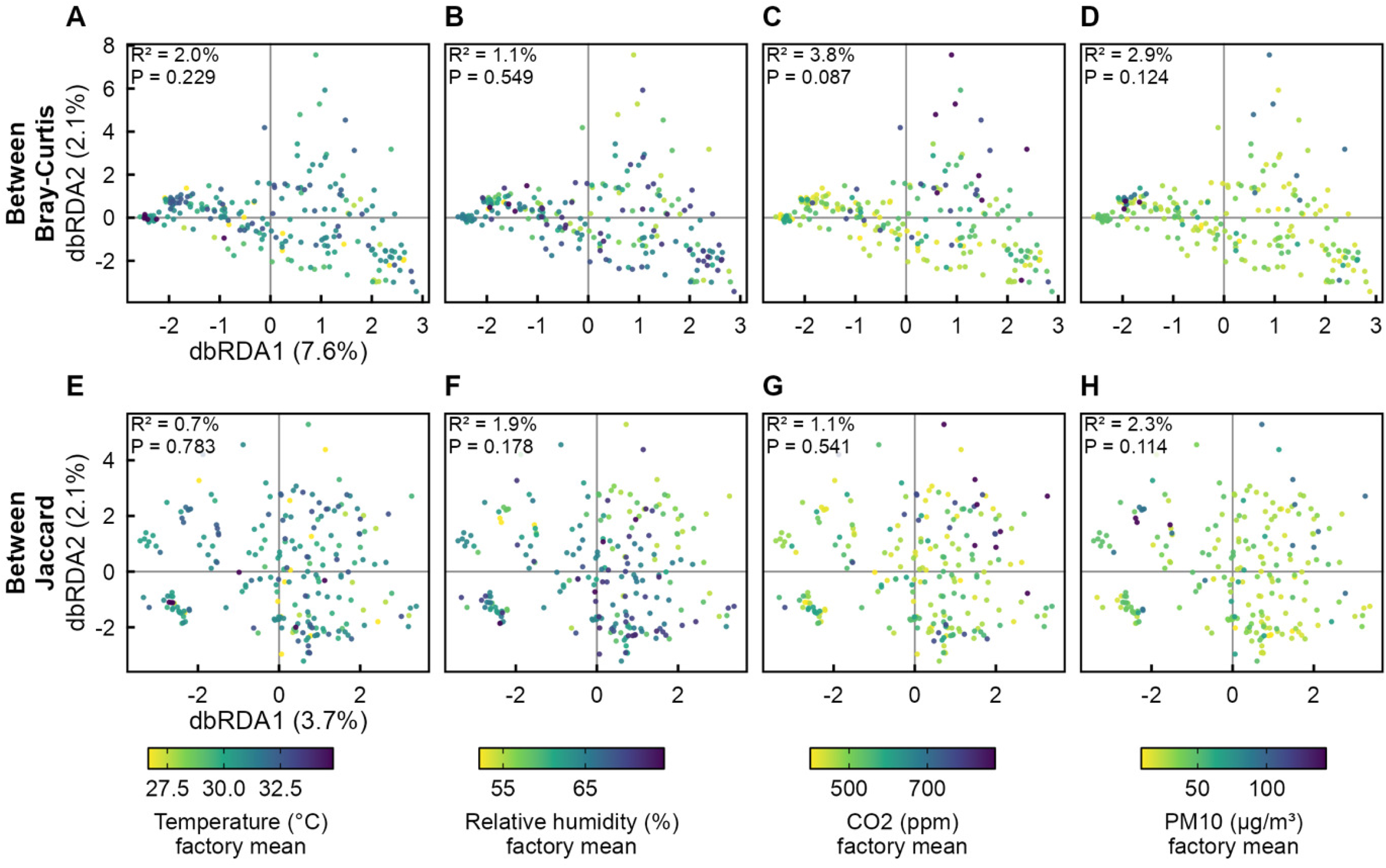

